# Engineered Retroviruses as Fluorescent Biological Reference Particles for Small Particle Flow Cytometry

**DOI:** 10.1101/614461

**Authors:** Vera A. Tang, Anna K. Fritzsche, Tyler M. Renner, Dylan Burger, Edwin van der Pol, Joanne A. Lannigan, George C. Brittain, Joshua A. Welsh, Jennifer C. Jones, Marc-André Langlois

## Abstract

There has been renewed interest in the use of flow cytometry for single particle phenotypic analysis of particles in the nanometer size-range such as viruses, organelles, bacteria and extracellular vesicles (EVs). However, many of these particles are smaller than 200 nm in diameter, which places them at the limit of detection for many commercial flow cytometers. The use of reference particles of diameter, fluorescence, and light-scattering properties akin to those of the small biological particles being studied is therefore imperative for accurate and reproducible data acquisition and reporting across different instruments and analytical technologies. We show here that an engineered murine leukemia virus (MLV) can act as a fluorescence reference particle for other small particles such as retroviruses and EVs. More specifically, we show that engineered MLV is a highly monodisperse enveloped particle that can act as a surrogate to demonstrate the various effects of antibody labeling on the physical properties of small biological particles in a similar diameter range.

---

Flow cytometry (FCM) analysis of viruses less than 500 nm in diameter has been reported since the late 70’s (1–3). Much of this foundational work required customized cytometer configurations, including high powered lasers, large collection angles, and very low sampling rates. Advances in the technology of modern cytometers now allow for some conventional commercial instruments to detect biological particles down to the 100 nm diameter range with minor to no modifications to default instrument configurations (4–14). However, several key challenges remain for small particle FCM and these include: variations in instrument configurations and detection capabilities across platforms and facilities, widely differing sample processing and labeling methods, and a lack of consensus for data reporting (15).

One of the major factors impeding these efforts for standardization is the paucity of available reference particles with fluorescence intensities relevant to that of biological samples (16). Reference particles are important for daily quality control of instrument performance, as well as internal positive controls for optimization of sample labeling protocols. Particles with low levels of fluorescence are needed to ensure optimal signal to noise resolution for dim signals. Biological reference particles have the advantage of possessing similar biochemical composition and antigen density levels, and can therefore act as suitable positive staining controls for antibody and dye labeling assays.

The development of calibration particles is also critical for the standardization of reference particle and sample data analysis and reporting in standard units. Although fluorescent reference particles in the form of polystyrene beads are commonly available, these are several microns in diameter and generally do not exhibit comparable fluorescence intensities (i.e., they are too bright) as EVs or even viruses. The calibration of fluorescence axes for small particle analysis using molecules of equivalent soluble fluorophore (MESF) units will require a set of populations that are smaller and dimmer than those currently available. Light scatter calibration using Mie modeling on the other hand requires homogeneous, well-characterized particles, with a variety of diameters and refractive indices (RIs) that are not necessarily the same as biological particles. Currently there are few sources of well-characterized light scatter reference materials for small particle FCM. Calibration particles and reference particles are therefore two distinct groups of materials that fulfill different roles with small particle standardization; the development of both, however, is required in the pursuit of standardized small particle FCM assays.

The murine leukemia virus (MLV) is symmetric and roughly spherical in shape, with a diameter of 124±14 nm as measured by electron cryo-microscopy (17). The specific strain used in this current study, Moloney MLV, is an ecotropic murine gammaretrovirus, meaning that it can only infect certain strains of susceptible mice(18). The viral envelope is primarily derived from the plasma membrane of infected cells, acquired during viral egress; a process that shares several common pathways with EV release into the extracellular medium (19,20). The precise and consistent stoichiometry involved in virion capsid assembly results in the release of particles that are monodisperse in structure. This is a critical and highly desirable feature, which distinguishes viruses from other biological reference particles. MLV naturally expresses on its surface host cell-derived markers, along with the viral envelope glycoprotein (Env). Env is expressed as a trimeric structure with a transmembrane domain (TM) and a surface (SU) antibody-accessible subunit (21,22). For most retroviruses, Env constitutes the only viral protein expressed on their surface. The number of Env trimeric structures, termed spikes, is a feature that has been characterized for several retroviral species. For example, while the human immunodeficiency virus type I (HIV-1) expresses approximately 14 - 21 spikes per particle, the simian immunodeficiency virus (SIV) was shown to have 73-98, the Rous sarcoma virus (RSV) ~82, and MLV ~100 (23–26).

For this study, we engineered a fluorescent MLV expressing superfolder GFP (sfGFP) as a fusion protein with Env (8,27). The fluorescence of Env-sfGFP was quantified using MESF beads (28,29). The unique features of viral homogeneity for both diameter and Env-sfGFP expression levels enabled the use of MLVsfGFP as a prototypic small vesicular particle to demonstrate quantification of fluorescence expression as a means to enumerate viral surface protein expression, as well as address pertinent questions regarding antibody labeling of small particles using Env-sfGFP as the target antigen. These include: 1) the relationship of fluorophore diameter and brightness to the resolution of small particle populations, 2) the impact of antibody labeling on diameter and RI, and 3) whether the use of multiple antibodies can impede optimal labeling and fluorescence intensities.

The use of this strain of MLV as a reference particle poses no biosafety concerns since they are ecotropic mouse viruses that are readily inactivated with formalin. They can also be lyophilized for stable storage and transport. The ability for them to be engineered to express surface epitopes of choice, fluorescent or otherwise, make these ideal controls for EV and virus immunophenotyping experiments. Based on these characteristics, we conclude that MLV particles exhibit essential features of a biological reference particle, and provide a much-needed tool for daily quality control, positive controls for select protein markers, and a simple method for evaluating cytometer sensitivity.

## MATERIALS AND METHODS

### MLV production

Generation of chronically infected NIH 3T3 cells and the production and preparation of MLV samples for flow cytometry analyses were described previously (8). MLVsfGFP was engineered from a glycogag-deficient MLV using overlapping primers to insert the sfGFP sequence into the proline-rich region of Env using a restriction-free cloning strategy, hence allowing for its surface expression as a chimera with the viral protein (27). Viruses were harvested from the supernatant of chronically infected NIH 3T3 mouse fibroblasts and directly analyzed by FCM. Briefly, for virus production, 2.5 × 10^6^ chronically infected cells were seeded into a 10-cm dish and cultured for 12 hrs. Cells were then washed to remove the serum-containing media and further cultured for 72 hrs in 10ml of phenol red-free DMEM (WISENT Inc.) supplemented with 10% (*ν/ ν*) EV-depleted fetal bovine serum. The cell supernatant was collected and passed through a 0.45 μm filter. The supernatant was then diluted with 0.1 μm-filtered PBS (WISENT Inc.) as required for analysis.

### Quantification of MLV particle concentration and coincidence detection

Viruses are produced at a constant rate by chronically infected cell lines. The concentration of virus in the supernatant from infected cells correlates directly with the number of infected cells seeded (Suppl. Fig. 1A). Particle concentration of viruses was determined based on virus-gated events using 1:1000 dilution, which usually yields a concentration of particles ~1-5×10^6^ particles/ml as determined by the Beckman Coulter CytoFLEX S with a sampling rate of 10µl/min. Volumetric counts obtained from the CytoFLEX were validated by NTA (Suppl. Fig. 1B and 1C). Serial dilutions of the MLV containing supernatants show consistent SSC and fluorescence intensities at dilutions below 1:500, with ≤1% electronic abort rate (Suppl. Fig. 1D to 1F).

**FIGURE 1.**
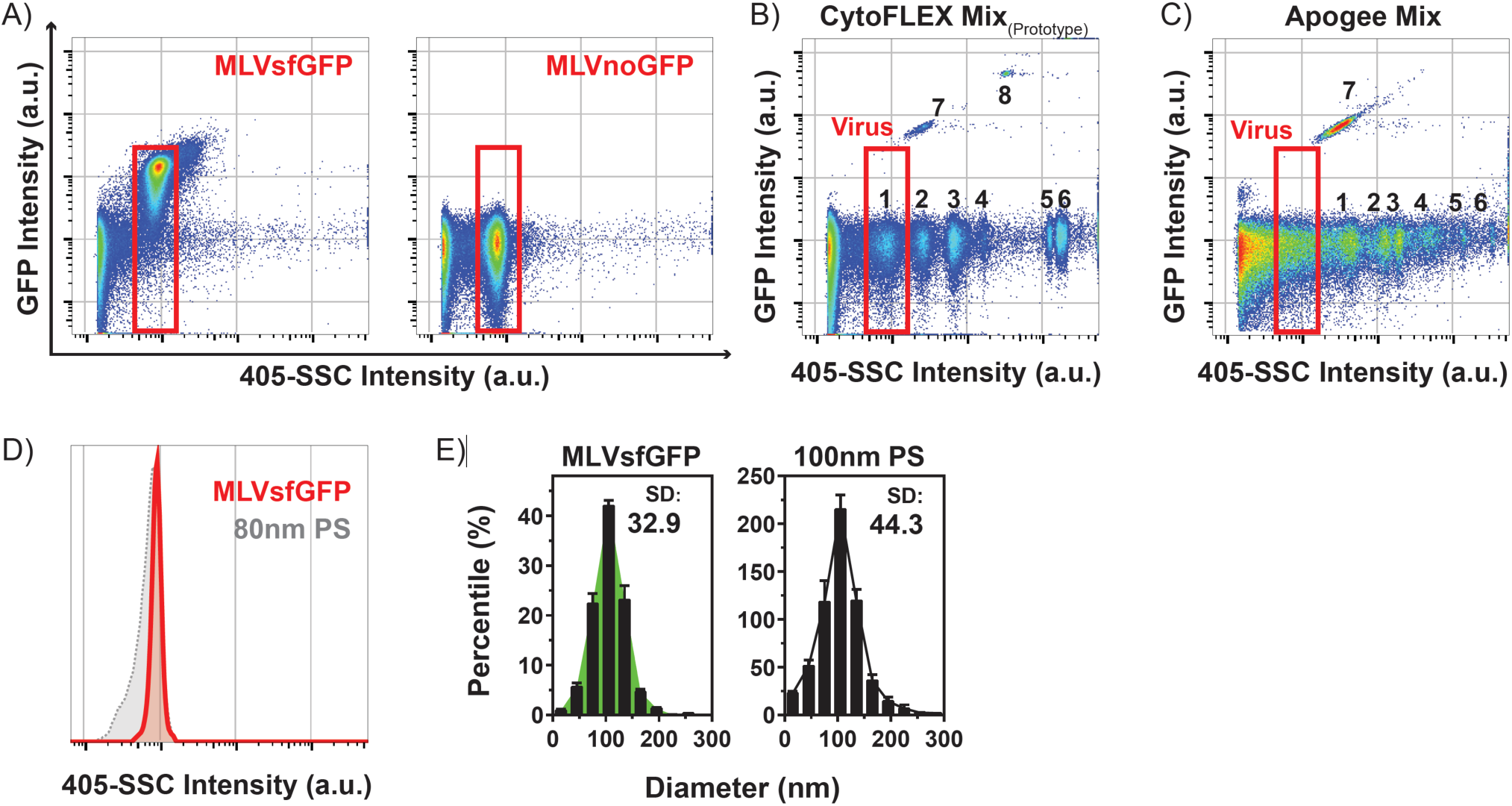
MLV virions are highly monodisperse. (A) MLVsfGFP and MLVnoGFP virions are a discrete population that can be resolved by 405-SSC intensity (a.u.) and green fluorescence intensity (a.u.). (B) Prototype CytoFLEX Sizing Mix (1: 80 nm PS, 2: 100 nm PS, 3: 214 nm Si, 4: 152nm PS, 5: 296 nm PS, 6: 1020 nm Si, 7: 100 nm PS fluorescent, 8: 196 nm PS fluorescent) and (C) ApogeeMix (1: 180 nm Si, 2: 240 nm Si, 3: 300 nm Si, 4: 590 nm Si, 5: 880 nm Si, 6: 1300 nm Si, 7: 110nm PS fluorescent) were analysed using the same settings as those for the MLV viruses. The “Virus” gate in red is the same from panel A. (D) Comparison of the 405-SSC intensity (a.u.) of MLVsfGFP and 80 nm polystyrene (PS) beads. (E) NTA on the diameter distribution of 100 nm PS beads and MLVsfGFP; SD: standard deviation.

### Flow cytometer set-up, beads, and data acquisition

Unless otherwise indicated, all samples were acquired on a Beckman Coulter CytoFLEX S with 4 lasers (405 nm, 488 nm, 561 nm, 640 nm), using 405 nm SSC-H (405-SSC-H) as the threshold parameter (threshold at 1400 a.u.). Detector gain for fluorescence and SSC detection were optimized using MLVsfGFP, with 0.1 µm-filtered PBS used as the background control for threshold determination. The gains for the respective detectors associated with the following spectral filters: 405-SSC, 405-450/50, 488-525/40, 561-580/30, and 640-670/30 were 1400, 1200, 3000, 1600, and 1200 a.u. respectively. A 405-SSC vs. time plot was used during acquisition to monitor, and ensure, consistency of the event rates. All samples were acquired for 1 min at a sampling rate of 10 μl/min. The sampling volume was validated by weight using the CytExpert volumetric calibration tool. The CytoFLEX Sizing Mix (prototype) (Beckman Coulter, Brea, CA) was analyzed undiluted and ApogeeMix (Apogee Flow Systems, Hemel Hempstead, UK) was diluted 1:5 with 0.1 µm-filtered PBS for analysis. FlowJo v.10 (FlowJo, LLC, Ashland, OR) was used for analysis of flow cytometry data.

### Imaging Flow Cytometry (IFC)

All MLV samples were acquired on a two camera ImageStreamX MKII (LuminexCorp.) according to the method previously described (12), with the modification of using the 405 nm laser (120 mW) for scatter measurements. Briefly, samples were acquired with 60X magnification, eGFP excitation with a 200 mW 488 nm laser, and scatter with the 405 nm laser described above. Emissions were collected for scatter in CH07 (bandpass 405-505nm) and CH02 for eGFP (bandpass 480-560 nm). All samples were acquired using the Inspire software and collected for a period of two minutes using a scatter acquisition gate that eliminated the speed beads (1µm polystyrene beads used for camera synchronization). Instrument sheath and sample dilution buffer was a 0.1 µm sterile filtered DPBS/Modified (HyClone cat. #SH30028.02). Buffer only controls were also run for the same amount of time to be sure that the same volumes were acquired as the samples. All virus samples were run in triplicate. 500 nm Si 7 peak FITC-MESF beads were also acquired using the same instrument settings as the virus samples. Data was processed using IDEAS 6.2 software (LuminexCorp) and FCS data files created for the scatter and GFP parameters and submitted for further analysis by the University of Ottawa Flow Cytometry and Virometry Core Facility.

### Fluorescence standardization and quantification using MESF beads

Calibration curves were generated using a linear fit by plotting the known MESF values vs. their respective fluorescence intensities in linear scale for each of the MESF bead sets used in these studies. The beads used were 500nm Si FITC-MESF (30), BD QuantiBrite PE (Lot 73318, BD Biosciences, Mississauga, ON), and Quantum-5 FITC MESF Beads (Bang Laboratories, Fishers, IN). Autofluorescence was measured using the blank bead population, and this was subtracted from the fluorescent-bead values. The uncertainties of the fluorescence values for each bead population was accounted for in the generation of the calibration curve and is represented as the standard error (SE), derived from the division of the standard of deviation (SD) by the square-root of counts obtained in each gated bead population. The linear fit of the calibration curve was weighted with the SEM of each bead population. The slope and intercept of each calibration curve for the 500 nm Si FITC-MESF and 7 µm PS FITC-MESF beads (Suppl. Fig. 2), was used to deduce the molecules of FITC equivalence for MLVsfGFP.

**FIGURE 2.**
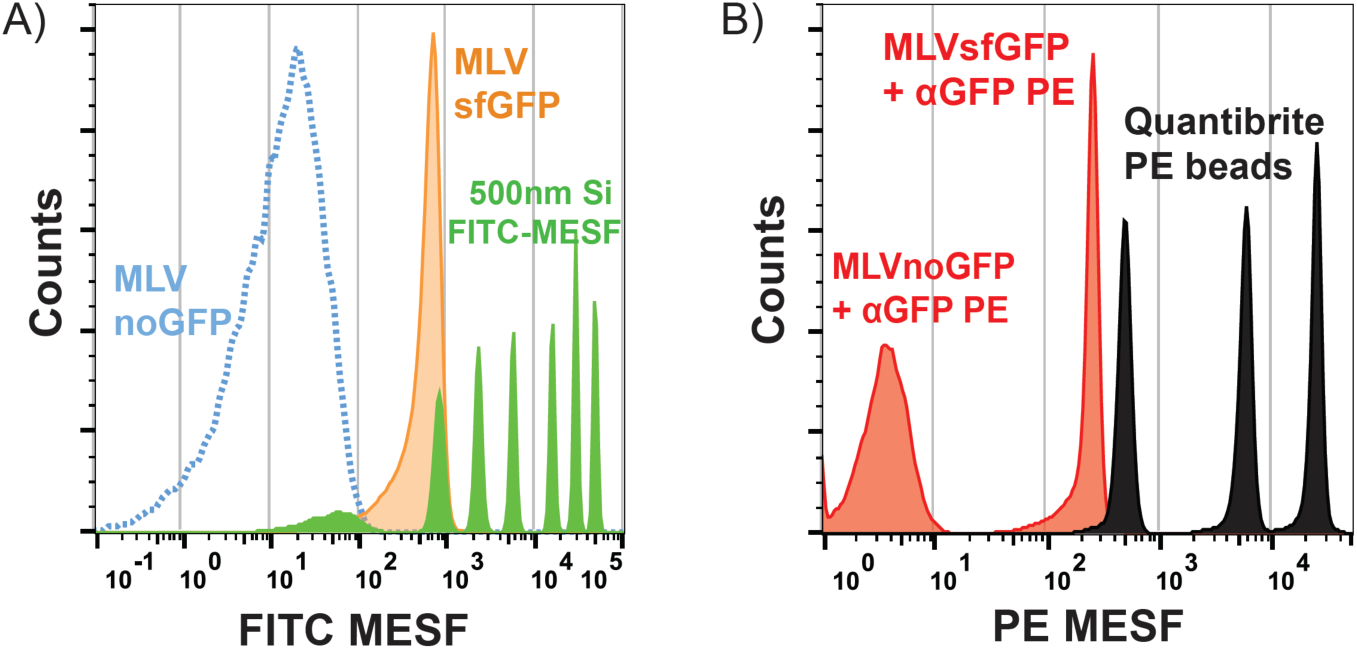
Fluorescence quantification of GFP expression on MLVsfGFP. (A) Green fluorescence from MLVsfGFP was quantified with 500nm Si FITC-MESF beads with MLVnoGFP as an autofluorescence control. Representative histogram overlay of MLVsfGFP and MLVnoGFP in FITC-MESF units with FITC-MESF beads (n=5). (B) Fluorescence quantification of anti-GFP PE labeled MLVsfGFP with QuantiBrite PE beads using MLVnoGFP as an internal control for non-specific labeling (n=3).

The virus population used for fluorescence quantification was identified based on its SSC and GFP fluorescence intensity and background fluorescence of the virus was subtracted using the fluorescence values of the gated MLVnoGFP. The mode of the sfGFP fluorescence intensities was used in determining the FITC-MESF value of MLVsfGFP. This statistic was chosen because it best represents the maximum of the unimodal distribution of our monodisperse virus population and is also the statistic most resistant to contributions from background noise events, which can be variable between day-to-day flow cytometer operations. The reported MFI and MESF values for MLVsfGFP was based on three separate experiments with a total of n=8 and n=9 samples. Calibration fits were produced using a C++ macro compiled with ROOT under the general public license (https://root.cern.ch/downloading-root). The slope and intercepts from the calibration fits were inputted into FlowJo to display the data as a derived parameter in terms of MESF units.

### Antibody labeling of MLV and MLV infected cells

For antibody labeling of MLV, the concentration of viral particles harvested from the supernatants of cells infected with MLVsfGFP and MLVnoGFP was adjusted to 10^9^ viral particles/ml for staining. Fluorophore-conjugated antibody aliquots were centrifuged at 17,000 × g for 10 min prior to use to reduce the presence of aggregates. For each antibody labeling reaction, 50 μl of virus supernatant was labeled with anti-GFP antibodies unconjugated or conjugated with PE, AF647 (clone FM264G, Bio Legend, San Diego, CA), or BV421 (clone 1A12-6-18), anti-mCD63-PE (clone NVG-2), anti-mCD81-PE or BV421 (clone Eat2), or anti-mCD9 PE (clone KMC8, BD Biosciences, Mississauga, ON) at the indicated concentrations for 1 hour at 37 °C in a total volume of 100 μl. For titration of all anti-GFP antibodies, 5 × 10^7^ MLVsfGFP viral particles were mixed with an equal number of MLVnoGFP particles, and a range of antibody staining concentrations from 0.0125µg/ml to 1.6µg/ml was tested for each anti-GFP conjugate. Unlabeled virus and antibody alone samples were run as controls for antibody labeling experiments. Labeled virus and controls were diluted 1:500 (~10^6^ particles/ml) for analysis in 0.1 μm-filtered PBS for analysis by FCM. For antibody labeling of MLV infected cells, 10^6^ cells were labeled with a concentration of 1 μg/ml of the same anti-tetraspanin antibodies used for MLV labeling in a 200 μl staining volume of 0.2% BSA-PBS for 20 min at 4°C. Excess antibody was removed by washing with 0.2% BSA-PBS. The SI was calculated for each anti-GFP conjugate at each concentration and the optimal staining concentrations associated with the highest SI value for anti-GFP PE, BV421, and AF647 were 0.2 µg/ml, 0.8 µg/ml, and 0.4 µg/ml respectively. The SI is defined as the difference of the MFI of the stained MLVsfGFP and MLVnoGFP divided by the standard of deviation of MLVnoGFP.

To assess the expression of cell-derived tetraspanins on MLV, MLVsfGFP was labeled with anti-mouse CD9, CD63, or CD81 antibodies conjugated with PE because this fluorophore was found to produce the highest SI. Gating strategy used to identify tetraspanin stained vs. negative particles is shown in Supplementary Figure 3 (panels A & B). Non-specific labeling with rat IgG-PE on MLV occurs at antibody concentrations greater than 1.6 µg/ml (Suppl. Fig. 4A). Lower concentrations of each antibody were also tested and confirmed that the optimal staining concentration (highest SI) was indeed 1.6 µg/ml (Suppl. Fig. 3C). Virus was identified by SSC intensity and gated to remove antibody aggregates using the antibody-only control samples (Suppl. Fig. 5A: red gates, and 5B: red events). PE and GFP intensities of anti-tetraspanin PE labeled fluorescent virus was converted to MESF using QuantiBrite PE and 500nm Si FITC MESF beads. QuantiBrite PE beads were chosen in this case because there are no commercially available small particle PE MESF beads.

**FIGURE 3.**
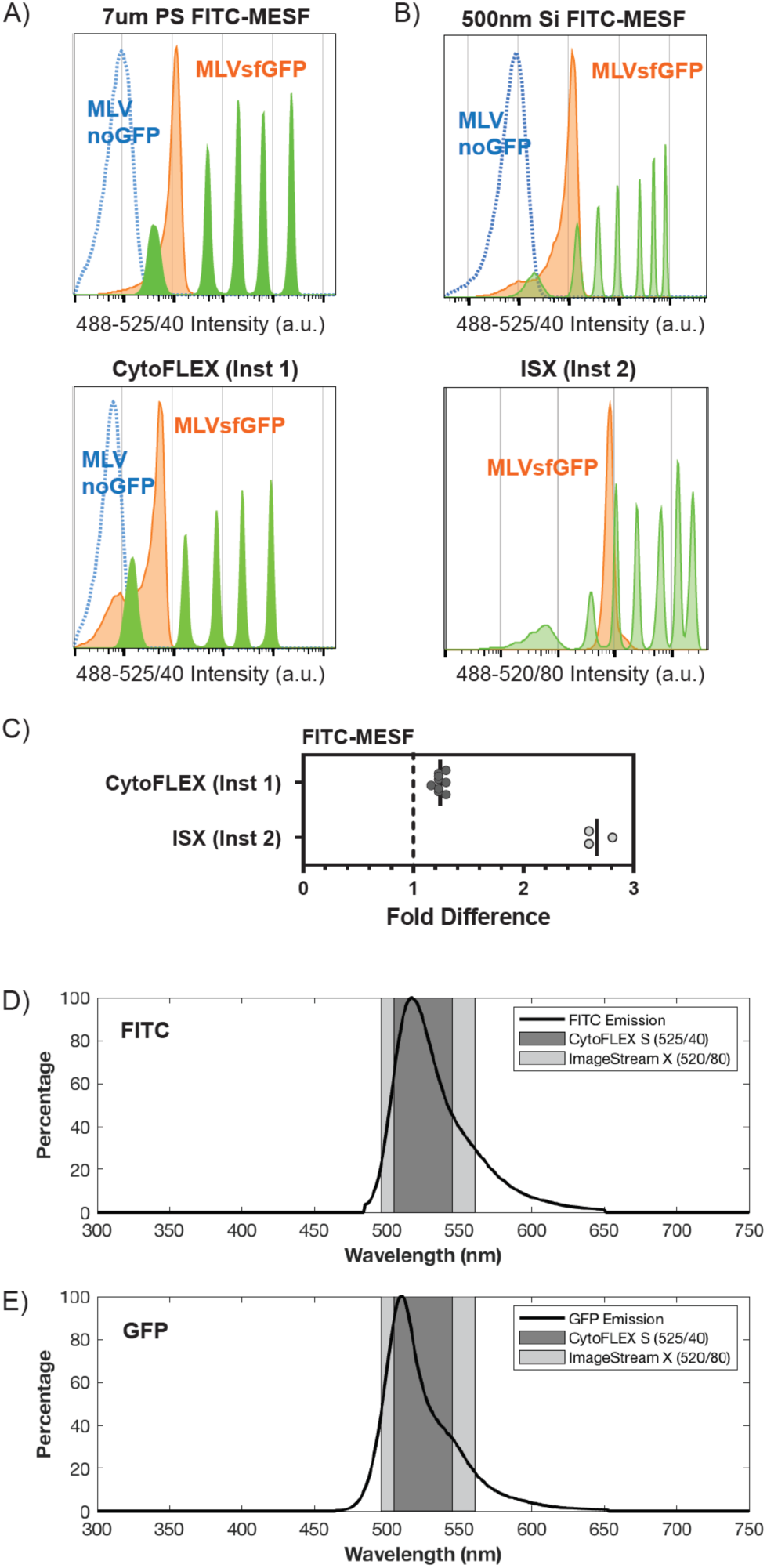
Cross-institution and cross-platform comparison of fluorescence intensity quantification of MLVsfGFP. (A) Comparative analysis of MLVsfGFP and MLVnoGFP viruses on two CytoFLEX S flow cytometers from two different institutions using 7µm PS FITC-MESF beads (top panel: uOttawa, bottom panel: Beckman Coulter (Inst1)). (B) Comparative analysis of MLVsfGFP and MLVnoGFP viruses on a Luminex ImageStream X (ISX) and a CytoFLEX S using 500nm Si FITC-MESF beads (top panel uOttawa CytoFLEX S, bottom panel uVirginia (Inst2) ISX). MLVnoGFP was not detected on the ISX. Data is displayed as fluorescence intensity. C) FITC-MESF values were calibrated for MLVsfGFP on both platforms and compared to values obtained with uOttawa CytoFLEX S, with uOttawa values set to “1” (dashed line). (D and E) Filter sets for the ISX and CytoFLEX S that were used for detection overlaid with the emission spectrums of FITC (D) and GFP (E); CytoFLEX S: n= 9; ISX; n = 3.

**FIGURE 4.**
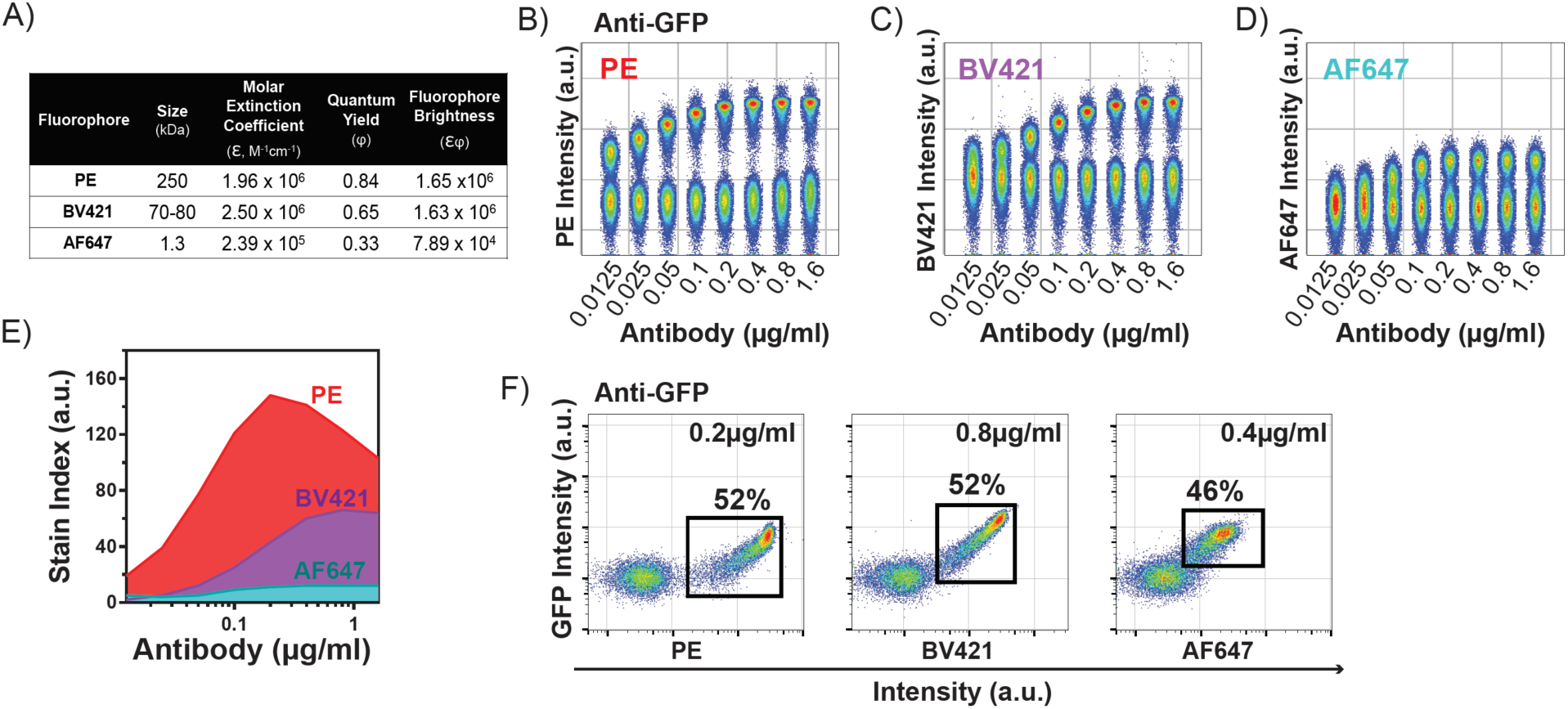
Evaluation of antibody-fluorophore conjugates for the detection of MLV. (A) Diameter and brightness information for PE, BV421, and AF647. Titration of anti-GFP (B) PE, (C) BV421, and (D) AF647 antibodies from 0.0125 µg/ml to 1.6 µg/ml, performed on a mixture of equal proportions of MLVnoGFP and MLVsfGFP virus particles. (E) The SI, displayed is a representative graph of n=6, was calculated for each antibody at each concentration to determine the optimal staining concentration. (F) Representative dot-plots showing the frequency of anti-GFP^+^ events labeled at optimal staining concentrations for each fluorophore conjugate.

**FIGURE 5.**
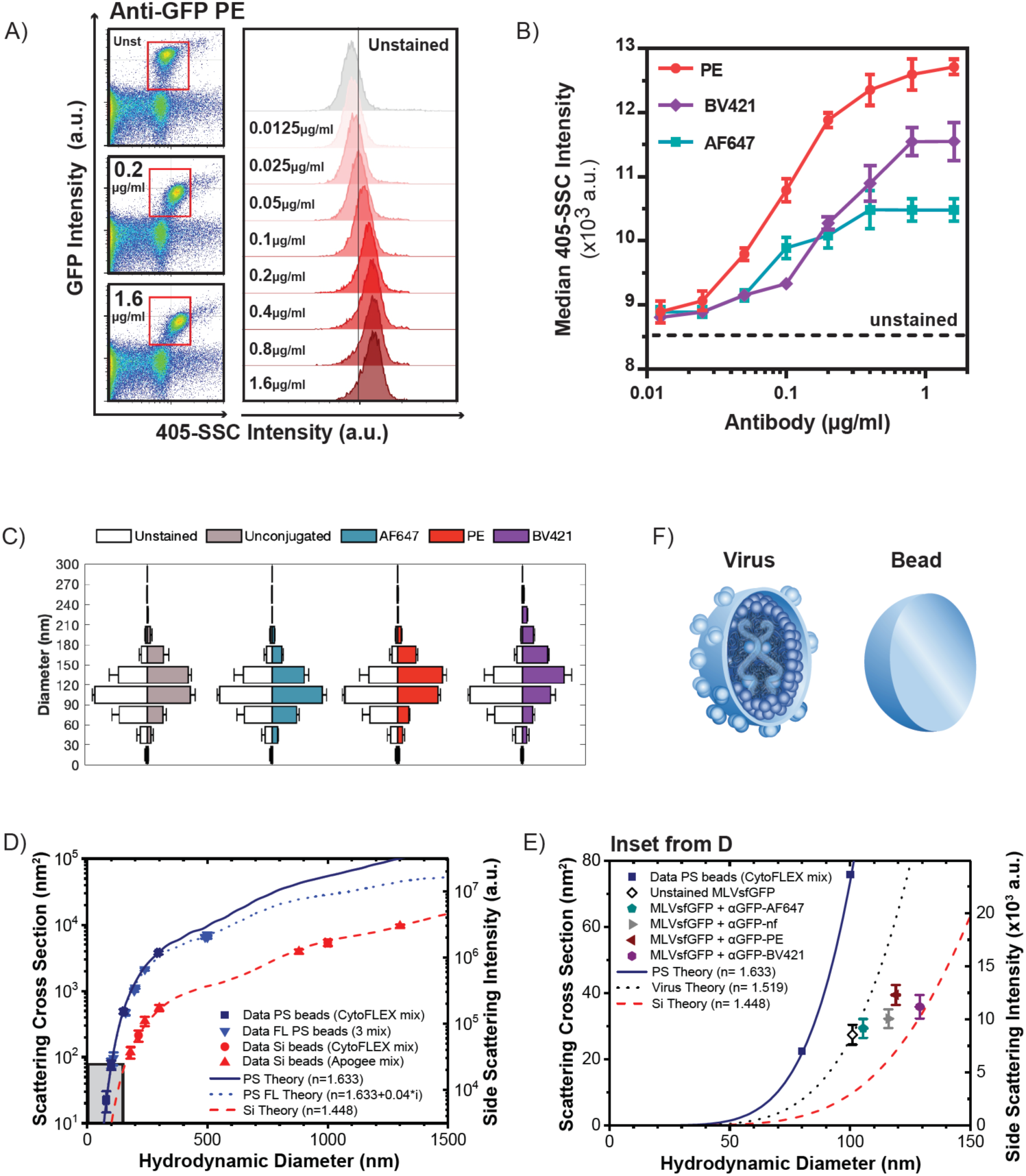
Antibody-fluorophore conjugates impact SSC intensity, hydrodynamic diameter, and effective refractive index of labeled viruses. (A) Antibody labeling of MLVsfGFP increases the SSC intensity of virus particles in a concentration-dependent manner. Representative scatter plots of MLVsfGFP (gated) mixed with MLVnoGFP that were unstained (Unst) or labeled with anti-GFP PE antibody at 0.2ug/ml and 1.6ug/ml (left panels). Histogram overlay of the gated GFP^+^ populations that were labeled with anti-GFP PE, for all antibody concentrations from 0.0125ug/ml to 1.6ug/ml (right panel). (B) SSC intensity of GFP^+^ viruses labeled with anti-GFP conjugated with PE, BV421, and AF647 at increasing antibody concentrations. Dashed line denotes SSC intensity of unstained MLVsfGFP. (C) NTA-measured diameter distribution of unstained MLVsfGFP separately compared to MLVsfGFP labeled with unconjugated anti-GPF, anti-GFP-PE, BV421, or AF647 at a concentration of 1.6µg/ml; n=3. (D) Mie-theory analysis for the calculation of the RIs for data points acquired using different silica (Si; red) and polystyrene (PS; blue) beads. The plot represents a correlation of the scattering cross section, hydrodynamic diameter, and SSC intensity of the virus particles. The gray-shaded box indicates the range where the MLV data points were acquired. (E) Gray-shaded box inset from (D), Mie-theory analysis of unstained MLVsfGFP and viruses labeled with various anti-GFP conjugated antibodies. The estimated effective RI of unlabeled virus is demonstrated as a dotted black line; n=3. (F) Cartoon to illustrate the complex and heterogeneous composition of a virus compared to the homogenous composition of a bead.

### Nanoparticle Tracking Analysis

NTA was carried out as previously described (8). Briefly, samples were diluted with 0.1 μm-filtered PBS and analysed using the ZetaView PMX110 Multiple Parameter Particle Tracking Analyzer (Particle Metrix, Meerbusch, Germany) in diameter mode using ZetaView software version 8.02.28. Camera gain: 938, Shutter: 70, Frame Rate 30 fps, Temperature 24.5, Brightness: 30. Videos were taken from all 11 camera positions.

### Light scatter modelling using Mie Scatter

Effective RI was approximated using the scatter-diameter curves based on the CytoFLEX S collection geometry as previously published (31). Briefly, instrument light scatter calibration was performed by fitting acquired 405 nm light scatter data to predicted values using the reported diameters and refractive indices of Apogee Mix beads (Apogee info) and NIST-traceable beads (Beckman Coulter, Brea, CA). These ranged in diameter from 80 nm polystyrene to 1300 nm silica (Fig. 1C).

### Data Sharing

All List-mode data files have been made available on *FlowRepository.org* in compliance MIFlowCyt Checklist item 4.1. Repository ID: http://flowrepository.org/id/FR-FCM-Z24Y.

## RESULTS

### MLV virions are monodisperse

The ecotropic Moloney MLV used for this study was modified to prevent the expression of the membrane-associated accessory glycogag protein (27). This alteration ensured that the only viral protein expressed on its surface is Env. This virus is termed MLVnoGFP in our study. MLVsfGFP consists on this same virus but with the insertion of sfGFP in the Env protein sequence. MLV virions were detected as a highly monodisperse population that could be resolved by side-scatter (SSC) intensity alone and further identified by GFP expression (Fig. 1A, red gates). Next, the 405-SSC intensity of the virus was compared to two types of sizing beads: CytoFLEX Sizing Mix (prototype, Beckman Coulter) (Fig. 1B) and ApogeeMix (Fig. 1C). The virus gate (red) from Figure 1A was superimposed to panels 1B and 1C to delineate where the fluorescent MLV population would appear with reference to the bead populations on our instrument system. MLV has a similar SSC intensity to 80 nm polystyrene beads (Fig. 1D). A comparison of the standard deviation (SD) in diameter distribution of MLVsfGFP and 100 nm polystyrene bead by nanoparticle tracking analysis (NTA) show a greater variability in diameter sizes in the beads versus virus, 44.3 and 32.9, respectively (Fig. 1E). This reflects the homogeneous and consistent stoichiometry of virus assembly, and suggests that formation of monodisperse MLVs is more consistent than the manufacturing methods currently used for production of NIST-traceable 100 nm polystyrene beads.

### Fluorescence quantification and enumeration of GFP molecule expression on MLVsfGFP

The fluorescence signal was analyzed from MLVsfGFP viral particles, using MLVnoGFP as the auto-fluorescence control and MESF calibration beads for fluorescence quantification (Fig. 2A). 500 nm silica spheres containing known MESF values of fluorescein isothiocyanate (500 nm Si FITC-MESF) (30) were used in lieu of GFP given that GFP-MESF beads in the relevant diameter and fluorescence-intensity range are currently not commercially available. The GFP intensity expressed by MLVsfGFP, quantified using the 500nm Si FITC-MESF beads, was found to be 637±3 FITC-equivalent molecules. Due to the mismatch of fluorophores between the FITC-MESF calibration beads and the MLVsfGFP, we could not report the fluorescence intensity of the virus in terms of GFP-MESF. Env-sfGFP expression was therefore quantified on MLV by an alternate method using anti-GFP-PE antibody labeling and fluorophore-matched QuantiBrite PE beads (Fig. 2B). A titration of the anti-GFP-PE antibody was performed to determine the concentration that would produce optimal labeling of Env-sfGFP using MLVnoGFP as an internal non-specific binding control (Fig. 4B and 4E). The brightest population of QuantiBrite PE beads were off-scale and only the first three populations were used. Anti-GFP-PE labeled MLVsfGFP had a PE-MESF value of 306±13, corresponding to 102 Env spikes. However, this quantification method also has potential limitations because it could underestimate the expression level of Env-sfGFP for several reasons: 1) inaccuracies associated with the use of MESF beads that are not calibrated for use with small particles, 2) quenching of PE molecules due to the proximity of target epitopes, 3) steric hindrance could prevent binding of all available epitopes, and 4) the bivalent nature of the antibody.

### Cross-institution and cross-platform assessment of fluorescence quantification

To compare the impact of 1) instrument variability, 2) user data acquisition variability, and 3) technological platform variations on the consistency of fluorescence quantification of the viruses, MLVsfGFP and MLVnoGFP viruses were sent to two different research institutions. The first institution operated a Beckman Coulter CytoFLEX S (Inst. 1) where virus fluorescence was quantified using 7 µm PS FITC-MESF beads (Fig. 3A). The second institution (Inst 2), operated a Luminex ImageStream X (ISX) where fluorescence quantifications were performed using 500 nm Si FITC-MESF beads (Fig. 3B). The values for FITC-MESF obtained on the sfGFP-expressing virus by Inst. 1 were very similar to our own, within a 0.25-fold difference, while Inst. 2 (ISX) produced values that were 2.6-fold higher (Fig. 3C). This apparent disparity was most likely due to differences in spectral filters between the two platforms. The width of the 525/40 bandpass filter used in the CytoFLEX S for collection of signal from FITC and GFP limits the collection of emitted photons to 62.7% and 59.2%, respectively (Fig. 3D and 3E). The wider filter on the ISX (520/80) is collecting 83.3% and 88.9% of photons emitted from FITC and GFP, respectively. This would suggest that the ISX was disproportionately collecting more signal from GFP than FITC (1.5-fold more compared to 1.3-fold), which could contribute to MLVsfGFP appearing brighter with respect to the FITC-MESF beads, highlighting a potential caveat of using mismatched fluorophores for fluorescence quantification.

### Antibody labeling of MLV surface antigens

FCM is the preferred method for immunophenotyping of cells. However, immunophenotyping of small particle populations, such as EVs and viruses, is inherently challenging due to low surface antigen abundance as a result of their restricted surface area. For optimal resolution of these populations, fluorophore selections are therefore limited to the brightest options, with minimal spectral spillover, thereby reducing the number of antigens that can be targeted in one antibody panel. In cells, fewer than 1000 molecules/cell is considered low antigen abundance (32). According to our own measurements, Env-sfGFP expression on MLV is potentially in the order of 10^2^ molecules (Fig. 2). Compared to a cell, this may seem low in total abundance, yet when integrated over surface area, this amount of antigen on a nanoparticle of ~100 nm in diameter actually translates to very high antigen density; the equivalent of several millions of molecules on a 10 μm cell. As a result, labeling of small particles with high antigen density could potentially present the challenge of steric hindrance issues that may occur for antibodies conjugated to larger fluorophores.

Many factors contribute to the number of photons detected by a flow cytometer from a fluorescently-labeled particle. These factors include: excitation wavelength, spectral filters, quantum efficiencies of detectors at increasing wavelengths, and the fluorophore to protein ratio (F:P ratio) of antibodies used to label the particles of interest. For the purposes of our study, instrument-specific considerations, such as excitation wavelength and spectral filters, are negated since analyses were performed on the same instrument. Avalanche photodiodes (APDs), the detectors used in the CytoFLEX S, also have a similar quantum efficiency over the range of visible light (400-800 nm) (33). The F:P ratio of conjugated antibodies, however, is a factor that should be considered, aside from the brightness, when choosing fluorophore conjugates since it is affected by the size of the fluorophore. Larger fluorophores such as PE typically have a 1:1 ratio due to steric hindrance, whereas smaller fluorophores could have a higher F:P ratio (34). Hence, a particle labeled with an antibody conjugated to the brightest fluorophore maybe not necessarily result in the greatest number of photons detected if the F:P ratio is low.

To assess the contributions of fluorophore size and brightness to resolving MLVsfGFP, an antibody against the high-density Env-sfGFP surface antigen was tested. Three different fluorophores that range in diameter and emission spectra, conjugated to an anti-GFP antibody were tested: PE, Brilliant Violet 421 (BV421), and Alexa Fluor 647 (AF647). The characteristics of each fluorophore, including brightness (εϕ) and size (kDa), are summarized in Figure 4A. PE is the largest and brightest of the three fluorophores, followed by BV421, and AF647. A titration was performed for the three conjugates of anti-GFP antibodies and the stain index (SI) was calculated for each (Fig. 4B to 4E). At optimal staining concentrations (highest SI), both the PE and BV421 conjugates identified an equivalent frequency of GFP^+^ viruses (52%), while the AF647 conjugate labeled slightly fewer GFP^+^ viruses than the other two fluorophore conjugates (46%) (Fig. 4F). As with labeling of cells, increase of the cell or particle concentration will decrease the SI of optimized antibody concentrations. We confirmed that at the optimal staining concentration of 0.2 µg/ml for anti-GFP PE, increasing the particle concentration of the sample does indeed decrease the SI, however this was only observed when particle concentrations increased by more than a factor of 2 (Suppl. Fig. 4B and 4C). We also observed that staining saturation is reached for the MLVsfGFP virus at a concentration of 1.6 µg/ml of the anti-GFP antibody (Suppl. Fig. 4A).

Although PE is a very bright fluorophore, one major caveat in using PE-conjugated antibodies for small particle FCM is its potential to form aggregates (35). In fact, PE^+^ particles were detected in samples containing only anti-GFP PE antibody which increased in number with rising concentrations of antibody used (Suppl. Fig. 5A). The majority of these PE^+^ aggregates were located in two populations (coloured events); one lower and the other higher than the labeled virus in SSC intensity (gray events) (Suppl. Fig. 5B). At the optimal staining concentration of 0.2 μg/ml for anti-GFP PE, the number of aggregates was negligible in comparison to the number of stained particles (Suppl. Fig. 5A & B). However, it is important to note that these samples were stained at 0.2 μg/ml, but then further diluted 1:500 for analysis, resulting in an actual antibody concentration of 0.4 ng/ml when analysed on the flow cytometer. Aggregates can also be seen with the anti-GPF BV421 conjugate, but not with the AF647 conjugate (Suppl. Fig. 5C and D).

### Antibody labeling of MLV modulates scatter intensity, hydrodynamic diameter, and the refractive index

During the analysis of our antibody-labeled MLVs in the previous section, we noted that the GFP^+^ virus populations increased in SSC intensity with increasing amounts of anti-GFP PE antibody (Fig. 5A; red gates). This increase in SSC was also observed with BV421 and to a lesser extent the AF647 conjugate (Fig. 5B). Conceptually, it is feasible that labeling with antibodies could significantly increase the apparent diameter of a small particle such as MLV. The diameter of an IgG antibody has been reported to range from 14 to 40 nm in diameter by 2 to 4 nm in height depending on the measurement method used (36,37). IgG conjugated with PE, which is 250 kDa and considered one of the larger fluorophores used in flow cytometry, has been reported to measure 60 nm in diameter by 5 nm in height by atomic force microscopy (35). To determine if the increase in SSC intensity is due to an increase in particle diameter, NTA was used to determine the hydrodynamic diameter of antibody-labeled MLVsfGFP at the over-saturating concentration of 1.6μg/ml. The median particle diameter and distribution were compared for unstrained MLVsfGFP and MLVsfGFP labeled with unconjugated anti-GFP, as well as PE, BV421, and AF647-conjugated antibodies (Fig. 5C). A scatter-modeling program based on Mie theory was used to calibrate the SSC intensity, to relate the SSC intensity to the measured hydrodynamic diameters of antibody-labeled and unlabeled MLVsfGFP determined with NTA (38), and to infer the RI. Data on the 405 nm scatter intensities acquired from polystyrene (RI=1.6333) and silica (RI=1.448) beads of known diameter (NIST-traceable) were used for calibration of our instrument (Fig. 1B). Our analyses showed high correlation (R^2^=0.9999) of acquired values (geometric symbols) with theoretical values (solid lines) down to 80 nm for PS (Fig. 5D). Theoretical lines represent Mie-theory simulations for materials of specific RIs with increasing particle diameter, scatter intensity, and scattering cross-section. Measured values for the diameters and scatter intensities of particles with the same RI are predicted to fall on the same lines as seen with the PS, fluorescent PS (FL PS), and Si beads (Fig. 5D). Figure 5E, generated from the gray inset in Figure 5D, depicts the collected data of antibody labeled MLV with respect to the RI values for PS (solid blue line) and Si (dashed red line).

Although individual MLV particles are not homogeneous in composition like a bead, their *effective RI* was calculated with this assumption to simplify the modeling of particles with multiple refractive indices due to mixed compositions (Fig. 5F). Here, the effective RI assumes the scattering intensity of each particle is related only to its refractive index and has no contributing extinction coefficient, as seen with fluorescent polystyrene beads (Fig. 5E). The dotted line, which passes through the unstained MLVsfGFP represents the effective RI of unstained virus (RI=1.519). The SSC intensity of antibody-labeled viruses falls below the iso-RI line of the unlabeled virus, indicating that labeled viruses have a lower effective RI than unlabeled virus. These results clearly show that antibody labeling can increase the diameter and, interestingly, reduces the effective RI, and therefore light scattering properties, of small particles. This may be related to the extinction coefficient of the fluorophore conjugated antibodies.

### Quantification of host cell-derived tetraspanins on MLV

Host-derived antigen expression on the surface of the virus by antibody labeling was next assessed to determine whether the observations from anti-Env-sfGFP labeling held true for other antigens on the surface of the virus. We chose to target cell-derived tetraspanins on the surface of MLV because these transmembrane glycoproteins are ubiquitously expressed as they contribute to fundamental processes of cellular trafficking (39). Tetraspanins CD9, CD63, and CD81 have been used as markers to identify subtypes of EVs due to their association with mechanisms of EV egress, such as the endosomal sorting complexes required for the transport (ESCRT) pathway (39). More specifically, these pathways have also been implicated in both cellular entry and egress of retroviruses (40–44).

The PE MESF of anti-tetraspanin-labeled viral particles were compared to show the relative expression levels of CD9, CD63, and CD81 on MLV (Fig. 6A-E). This comparison is possible because, in contrast to other fluorophores, only one PE molecule is likely to be conjugated per IgG due to its large diameter (34). CD81 was most abundantly expressed on MLVsfGFP with a median PE MESF of 18.7±0.2, followed by CD63 with 13.1±0.3 PE MESF, and CD9 with 6.4±0.02 PE MESF. It is important to note that these values should be taken as a measure of relative tetraspanin abundance between the three types, and not as actual molecules of tetraspanins expressed per virus, since QuantiBrite PE beads were not intended for use with such dimly expressed antigens and so are not accurately calibrated for this purpose (45). It is unclear whether CD9 expression was actually present on MLVsfGFP since the signal was similar to unstained virus (3.6±0.02 PE MESF) and could potentially be the result of non-specific labeling. However, CD9 was confirmed to be expressed on the cells producing MLVsfGFP, therefore it is possible that MLVsfGFP indeed express CD9 at very low levels, below the detection limit of our flow cytometer (Suppl. Fig. 3D).

**FIGURE 6.**
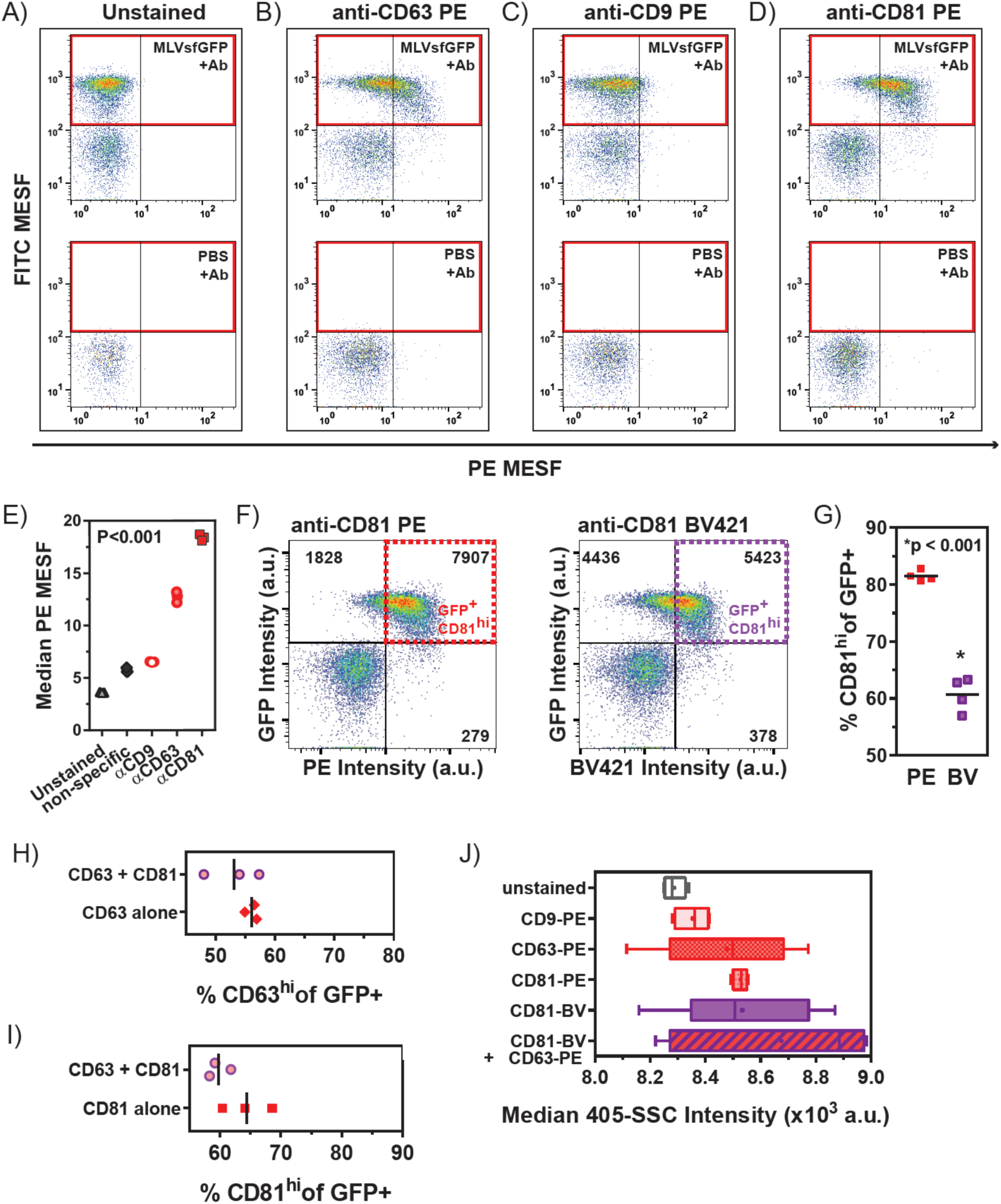
Phenotypic analysis of cell-derived tetraspanins expressed on the surface of MLV virions. (A) Unstained MLVsfGFP was compared to virus labeled with (B) anti-CD63-PE, (C) anti-CD9-PE, and (D) anti-CD81-PE at a concentration of 1.6 μg/ml of antibody per sample. QuantiBrite PE beads and 500nm Si FITC MESF were used to convert fluorescence intensity to PE and FITC MESF. (E) Median PE-MESF values of anti-tetraspanin labeled MLVsfGFP (n=3), 1-way ANOVA, P<0.001. (F) Gating strategy for identification of GFP^+^CD81^hi^ population from MLVsfGFP labeled with anti-CD81 BV421 and anti-CD81 BV421. (G) Comparison of GFP^+^CD81^hi^ events from anti-CD81 PE and anti-CD81 BV421 labeling, n=3, Unpaired t-Test, p<0.001. (H) Comparison of the frequency of CD63^hi^GFP^+^ virus in single-(anti-CD63 alone) vs. double-labeled (anti-CD63 + anti-CD81) viruses, Unpaired t-Test, non-significant, p = 0.34. (I) Frequency of CD81^hi^GFP^+^ virus in single-(anti-CD81 alone) vs. double-labeled (anti-CD63 + anti-CD81) viruses, Unpaired t-Test, non-significant, p = 0.15. (J) SSC intensities (405-SSC) for anti-tetraspanin labeled viruses, 1-way ANOVA, non-significant, P = 0.16.

The level of staining produced by the same anti-CD81 antibody conjugated to PE vs. BV421 was compared (Fig. 6F). Labeling of MLVsfGFP with anti-GFP-PE resulted in a higher SI than with the BV421 conjugate, although both equally resolved the MLVsfGFP population at optimal staining concentrations (Fig. 4B, 4C, and 4F). At optimal staining concentrations, labeling MLVsfGFP with anti-CD81-PE resulted in approximately 20% higher frequency of CD81^+^GFP^+^ viruses than anti-CD81-BV421 (Fig. 6G). The resolution of CD81 expression, an antigen expressed at lower levels than Env-sfGFP, benefited significantly from the use of a brighter fluorophore.

To assess whether double-labeling, the targeting of two different antigens with two different antibodies, would result in reduced staining for each individual antigen due to possible steric hindrance between the fluorophore-conjugated antibodies. A comparison was made between the percent of resulting CD81^+^GFP^+^ and CD63^+^GFP^+^ viruses (based on the gating strategy used in Fig. 6F; dashed gates) when the virus was labeled with anti-CD63 PE and anti-CD81 BV421 individually or with both antibodies together. There was no significant difference between the numbers of CD81^+^MLVsfGFP or CD63^+^MLVsfGFP events, or in the percentage of CD81^+^ or CD63^+^ GFP^+^ events, obtained using single versus double-labeling (Fig. 6H and 6I). This suggests that steric hindrance did not affect in this case the individual binding of two antibodies targeting distinct antigens.

SSC intensities for single and double-labeled virus populations (Fig. 6J) were compared to determine if fluorescence labeling of lower-density antigens would similarly impact scatter intensity and, thus, the apparent diameter and RI. Although there was an appearance of a correlation between the highest SSC intensities and the highest degree of labeling (CD81+CD63>CD81>CD63>CD9), these values were not statistically different from those of the unlabeled virus. Therefore, these observations suggest that the labeling of low-density antigens with fluorophore-conjugated antibodies does not significantly alter the scatter intensities of small particles.

## DISCUSSION

Current challenges for the analysis of small biological particles by flow cytometry are multi-faceted. To reliably achieve single-molecule resolution, technological advancements are needed to further improve instrument sensitivity. Development of brighter and smaller fluorophores is required for multi-parameter analyses of small particles where surface area is highly restricted. But more urgently, different types of reference particles with low fluorescence levels relevant to small particles are also needed for: 1) positive controls for stainings (antibodies and dyes), and 2) instrument calibration to allow standardized data reporting across instruments and technological platforms.

At present, few references particles are available for small particle FCM. Many FCM reference beads, such as compensation beads, are made mostly of polystyrene and exhibit fluorescence and autofluorescence intensity levels that are much higher than those attainable with small biological particles. Chemical conjugation of biological molecules to synthetic beads for use as positive controls can be technically challenging and often result in very high levels of expression that are biologically irrelevant. Currently available calibration beads are also too large and too bright for accurate small particle fluorescence standardization using MESF axes calibration. Biological particles, such as retroviruses, on the other hand, have long been adapted for use as vectors to safely express proteins of interest in cells. MLV particles are small, monodisperse, and have minimal autofluorescence. The potential of MLVs for use as fluorescence calibration particles in FCM is obvious because they can be engineered or labeled to have multiple levels of fluorescence expression. We showed here, with titrating levels of anti-GFP antibody, that MLVsfGFP can be easily labeled with different intensities of fluorescence.

In this study, MLV was engineered to express sfGFP in fusion with the viral surface glycoprotein, Env. These virus particles were used to showcase the importance of FCM best practices, such as antibody titration and fluorophore selection, when conducting immunophenotyping assays on small biological particles. Additionally, we demonstrated that when a highly-expressed surface antigen on a small particle is labeled with an excess of fluorophore-conjugated antibodies, these can change the physical properties, including the diameter and effective RI of that small particle. Additionally, this study emphasized the importance of fluorescence standardization with matching fluorophores to compare data between different flow cytometry platforms. Taken together, our observations on antibody labeling using MLV as a prototypical small particle, enabled us to identify and address specific challenges relevant to the antibody-labeling of small biological particles. These observations were only made possible due to the stringent uniformity in diameter and fluorescence, and high viral surface antigen expression on MLV particles. These critical features decisively qualify MLV as a candidate biological reference particle for the FCM analysis of other enveloped viruses and small biological particles such as EVs.

## ACKNOWLEDGEMENTS

The authors would especially like to thank members of the Beckman Coulter technical support team, Dominic Therrien and Marc Simard, for valuable assistance throughout this study. We would like to acknowledge Sergei Gulnik and Maria Gentile of the Beckman Coulter Research and Marketing Teams for their helpful discussions. We would like to thank Christian Ouellet for providing the C++ scripts for MESF calibration. V.A.T. is an International Society for Advancement of Cytometry (ISAC) Shared Resource Lab Emerging Leader. T.M.R. holds a Queen Elizabeth II Graduate Scholarship in Science and Technology (QEII-GSST). E.v.d.P. was supported by the Netherlands Organization for Scientific Research-Domain Applied and Engineering Sciences (NWO-TTW), research programs VENI 15924. M.-A.L. holds a Canada Research Chair in Molecular Virology and Intrinsic Immunity. This work was supported by a research and development grant from the University of Ottawa Faculty of Medicine to the FCV Core Facility, and by a Discovery Grant and an Idea to Innovation (I2I) Grant by the Natural Sciences and Engineering Research Council of Canada (NSERC) to M.-A.L.

## SUPPLEMENTARY MATERIAL

**Supplementary Figure 1.**
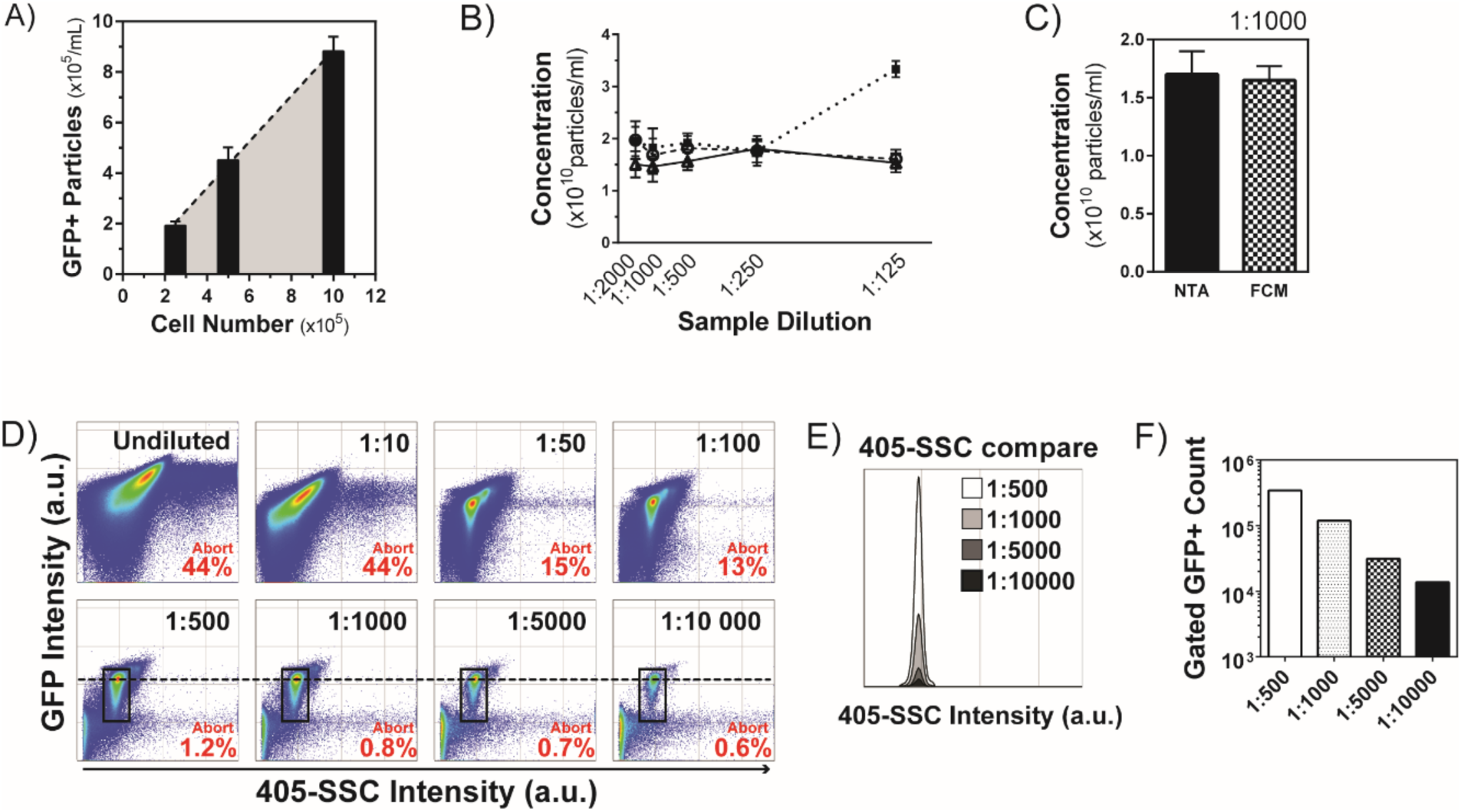
Coincidence detection and determination of virus particle concentration through serial dilutions using FCM and NTA. A) Analysis of equivalent volumes of supernatants collected from an MLV infected cell line correlating seeding densities of cells to the amount of virus produced; n=6. B) Calculated particle concentration of MLVsfGFP in undiluted supernatant based on concentrations measured by NTA in serial dilutions (n=3). C) The calculated concentration of undiluted MLVsfGFP supernatant based on measurements determined by NTA and FCM using samples diluted 1:1000 (n=3). D) Flow cytometry analysis of MLV dilutions showing the abort rates and increase in measured GFP fluorescence intensity and 405-SSC intensity of MLVsfGFP at the highest concentrations. E) An overlay of the events from the highest dilutions to compare 405-SSC scatter intensities. F) Linear correlation of GFP+ events (gated in (Fig. 1A) with dilution factor.

**Supplementary Figure 2.**
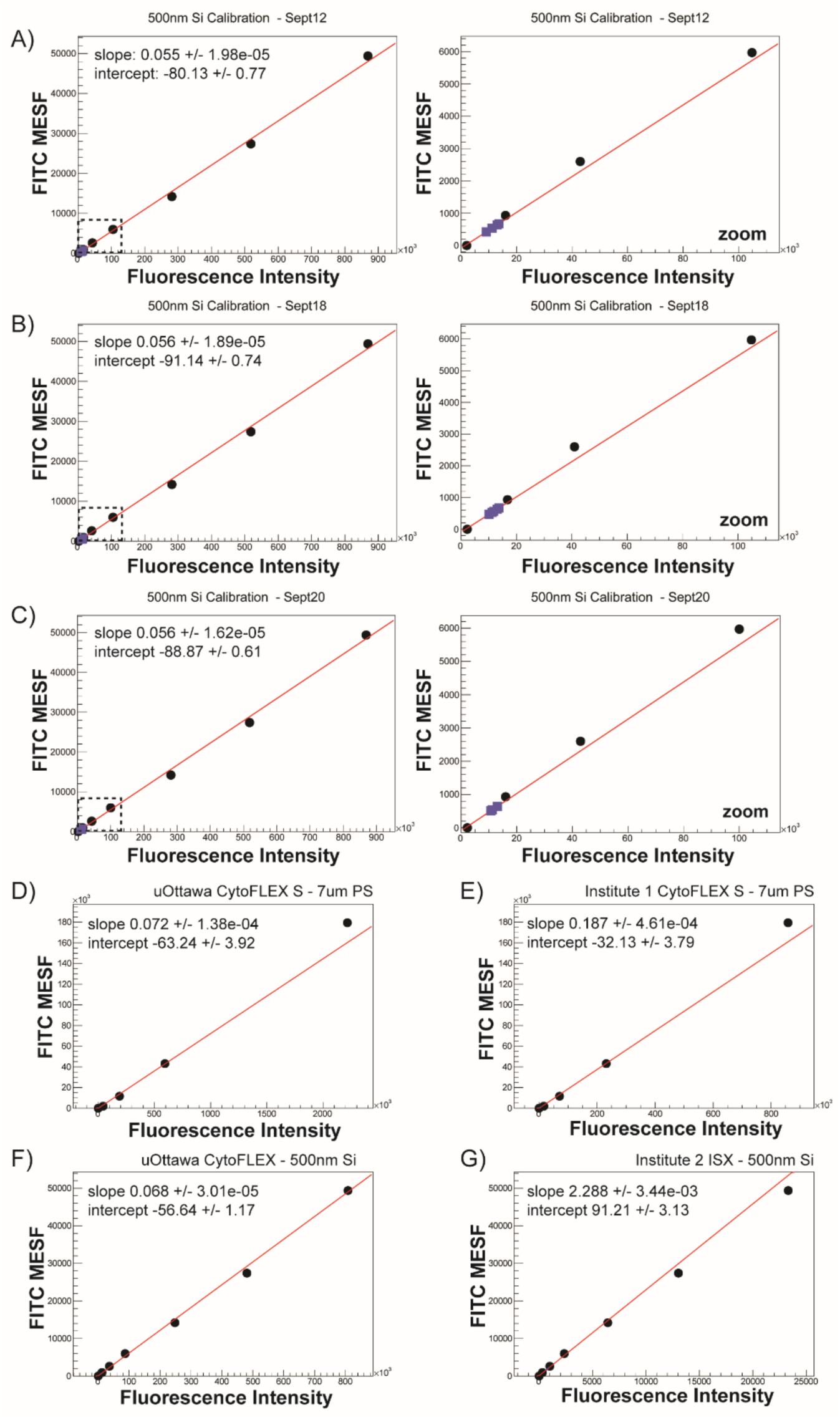
Calibration Curves for MESF calculations for Figures 2 & 3. (A to C) Calibration curves for 500nm Si MESF beads for data collected on 3 separate dates for MESF values summarized in Figure 2G. (D) Calibration curve for uOttawa CytoFLEX S using 7 μm PS FITC MESF Beads. (E) Calibration curve for Institute 1 CytoFLEX S using 7 μm PS FITC MESF Beads. (F) Calibration curve for uOttawa CytoFLEX S using 500 nm Si FITC MESF Beads. (G) Calibration curve from Institute 2 ISX using 500 nm Si FITC MESF Beads.

**Supplementary Figure 3.**
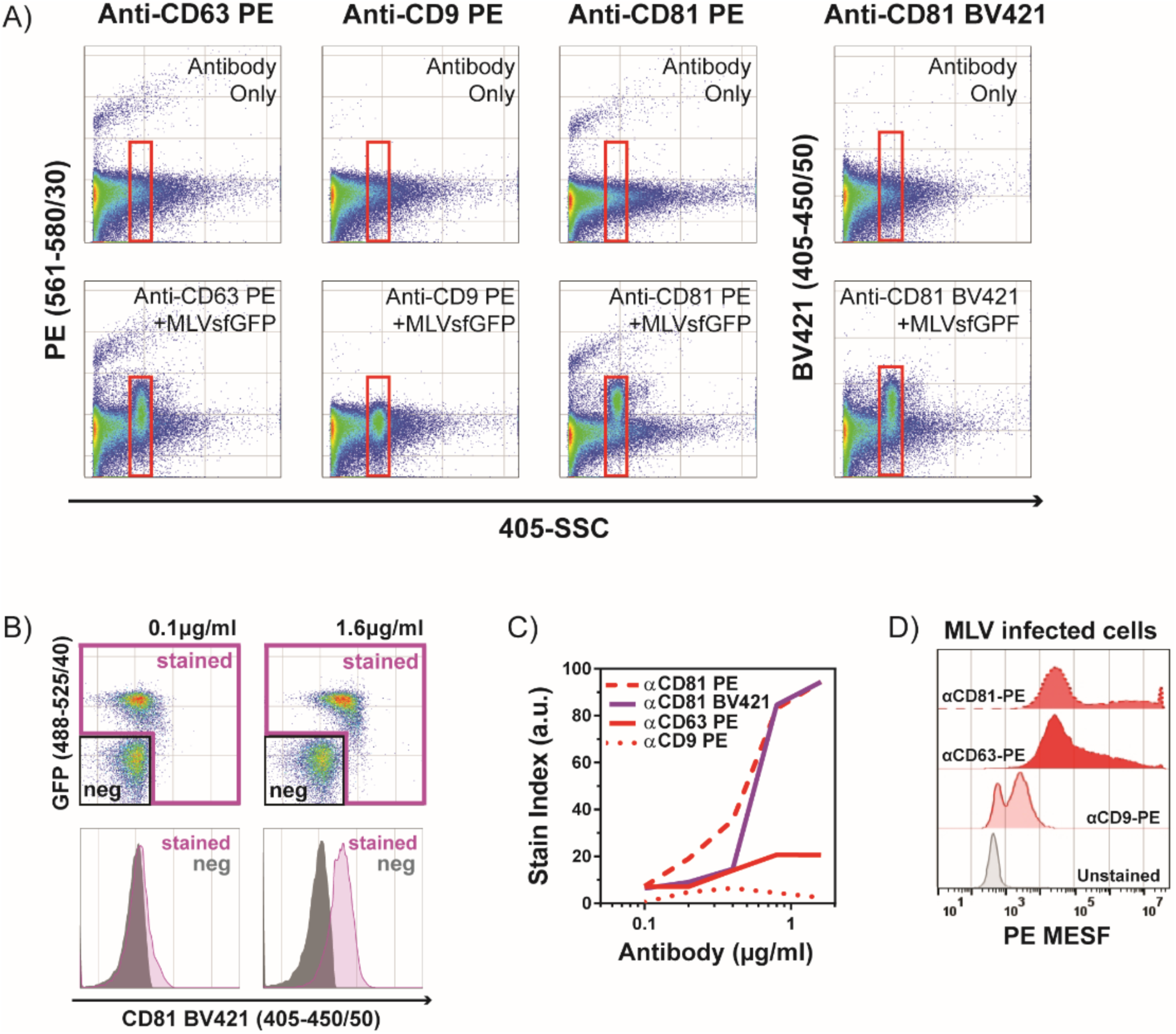
Anti-tetraspanin labeling of MLVsfGFP and MLVsfGFP infected cells. (A) Gating strategy for virus events to remove antibody aggregates using antibody only controls. All events are displayed as PE Intensity vs. 405-SSC Intensity for anti-mouse PE conjugates for CD9, CD63, and CD81. (B) Gated events from (A) are displayed as GFP vs. PE Intensity. These events were then calibrated to be displayed as FITC MESF vs PE MESF in Figure 6. (B) Gating strategy used to identify stained and negative populations used to calculate SI. (C) SI for anti-CD81-PE, anti-CD81-BV421, anti-CD63-PE, and anti-CD9-PE at concentrations from 0.1 μg/ml to 1.6 μg/ml. D) Anti-tetraspanin labeling of chronically infected producer cells for MLVsfGFP, representative histogram of n=3.

**Supplementary Figure 4.**
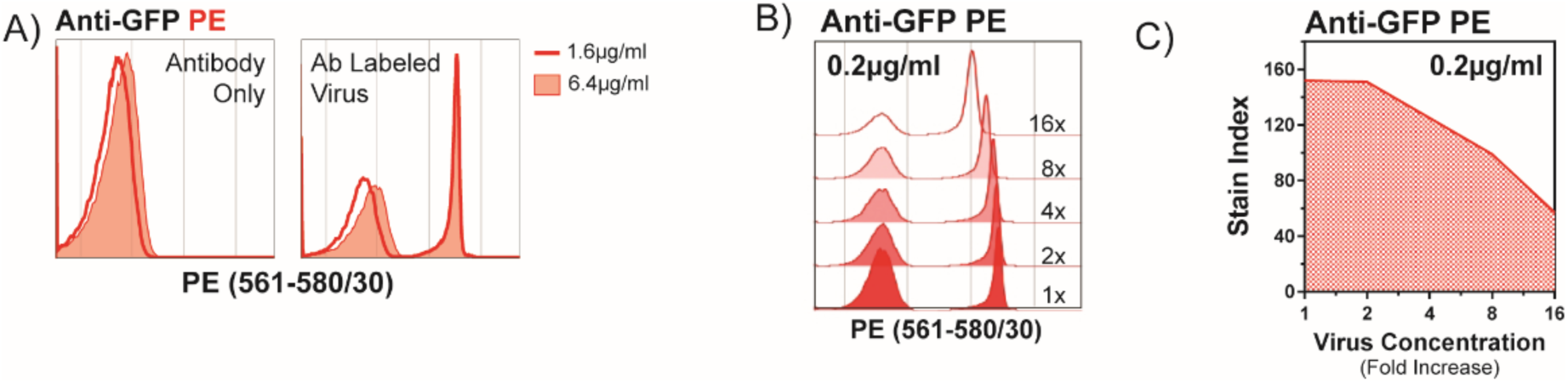
Effect of changes in particle and antibody concentration on the stain index of antibody labeled MLVsfGFP. (A) Histogram overlay of MLVsfGFP labeled with anti-GFP-PE at increasing virus concentrations while maintaining staining concentration of 0.2 µg/ml. 1x is the original virus concentration used to obtained optimal stain index at 0.2 µg/ml. (B) Stain index calculated from (A). (C) Histogram overlay of anti-GFP-PE antibody-alone at 1.6 µg/ml and 6.4 µg/ml (left panel) and anti-GFP-PE labeled MLVsfGFP + MLVnoGFP (right panel) showing staining of the MLVsfGFP population is saturated at 1.6 µg/ml.

**Supplementary Figure 5.**
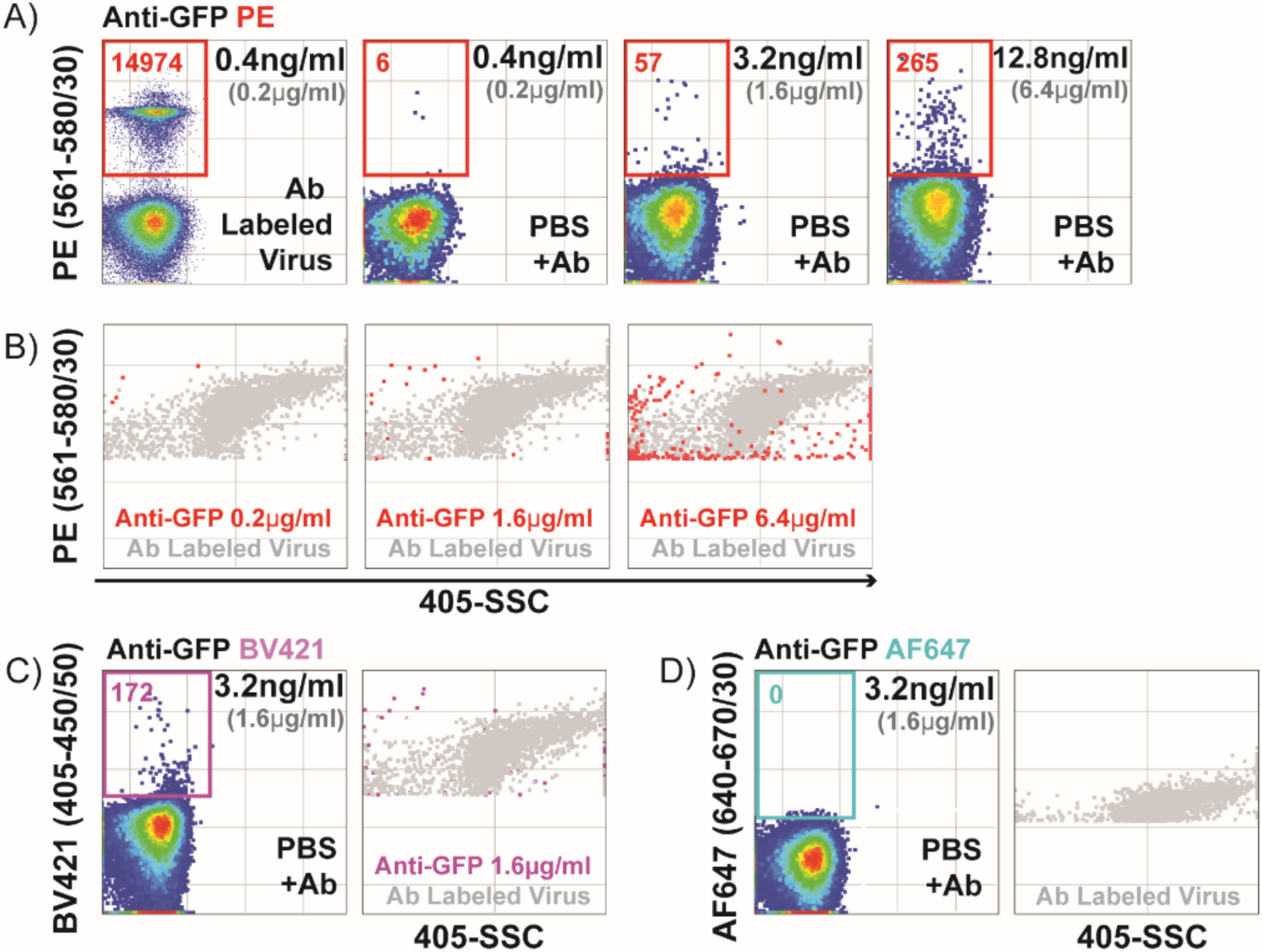
Detection of antibody aggregates in anti-GFP conjugates. (A) Dilutions of anti-GFP-PE antibody alone were analyzed at 0.2μg/ml (optimal staining concentration), 1.6μg/ml and 6.4μg/ml. The first panel on the left denotes MLVsfGFP stained at a concentration of 0.2µg/ml. Concentrations in black indicate the actual concentration of antibody as it is diluted for analysis on the cytometer. Values in red are PE+ event counts within the red gate. (B-D) Overlays of MLVsfGFP labeled at optimal staining concentration (gray events) with increasing concentrations of (B) anti-GFP-PE; (C) 1.6μg/ml of anti-GFP-BV421; and (D) anti-GFP-AF647. Representative plots for three independent experiments are shown.

